# Knockout of ribosomal protein RpmJ leads to zinc resistance in *Escherichia coli*

**DOI:** 10.1101/2022.10.24.513496

**Authors:** Riko Shirakawa, Kazuya Ishikawa, Kazuyuki Furuta, Chikara Kaito

## Abstract

Zinc is an essential metal for cells, but excess amounts are toxic. Other than by regulating the intracellular zinc concentration by zinc uptake or efflux, the mechanisms underlying bacterial resistance to excess zinc are unknown. In the present study, we searched for zinc-resistant mutant strains from the Keio collection, a gene knockout library of *Escherichia coli*, a model gram-negative bacteria. We found that knockout mutant of RpmJ, a 50S ribosomal protein, exhibited zinc resistance. The *rpmJ* mutant was sensitive to protein synthesis inhibitors and had altered translation fidelity, indicating ribosomal dysfunction. In the *rpmJ* mutant, the intracellular zinc concentration was decreased under excess zinc conditions. RNA sequence analysis revealed that *rpmJ* knockout decreased the expression of synthetic genes for iron-sulfur cluster proteins, which are toxic targets for zinc. These findings suggest that knocking out RpmJ causes zinc resistance by decreasing zinc targets and lowering the intracellular zinc concentration. Knockouts of other ribosomal proteins, including RplA, RpmE, RpmI, and RpsT, also led to a zinc-resistant phenotype, suggesting that deletion of ribosomal proteins is closely related to zinc resistance.

## Introduction

Zinc is an essential metal for organisms. Approximately 5% to 6% of total proteins in bacteria are zinc-binding proteins [1]. Zinc acts as a cofactor for enzyme activity and protein structure folding. On the other hand, excess zinc is toxic to cells by destroying [4Fe-4S] clusters of dehydratases and releasing free irons [2]. Iron, a metal with high redox potential, produces reactive oxygen species by the Fenton-reaction and impairs cell growth [2–4].

Bacteria must maintain a strict intracellular zinc concentration to reserve a necessary amount of zinc while avoiding toxicity from excess zinc. Four main zinc transporters have been identified in *Escherichia coli*. ZnuABC [2], a high-affinity ABC transporter, and ZupT [5], a ZIP family transporter, are responsible for zinc uptake. Under zinc-deficient conditions, the expression of ZnuABC is upregulated by relieving the transcriptional repressor Zur, a homolog of Fur [6]. ZntA, a P-type ATPase transporter [7, 8], and ZitB, a cation diffusion facilitator family transporter, mediate zinc efflux [9]. Under excess zinc conditions, the transcription factor ZntR upregulates the expression of ZntA [10, 11]. Other than the zinc efflux and uptake systems, little is currently known about the factors involved in zinc resistance. In the present study, we aimed to identify the genetic factors responsible for zinc resistance utilizing a gene knockout mutant *E. coli* library. We found that knockout of the 50S ribosomal protein RpmJ conferred zinc resistance. The *E. coli* ribosome contains 54 proteins, of which RpmJ is 1 of 8 nonessential ribosomal proteins. RpmJ is the smallest 50S ribosomal protein with only 38 amino acids [12], and is involved in 23S rRNA folding [13]. In this study, we investigated the mechanism of zinc resistance in the *rpmJ* knockout mutant by analyzing gene expression and the intracellular zinc concentration.

## Results

### Knockout of *rpmJ* causes zinc resistance

In this study, we searched a gene knockout mutant library for gene knockout mutants that grew on Luria broth (LB) agar plates containing zinc to identify genetic mutations that confer zinc resistance to *E. coli*. Four zinc-resistant mutant strains were identified (**Table 1**) with the *rpmJ* mutant exhibiting the strongest zinc-resistant phenotype. The other 3 mutant strains were *pitA*, *rimP*, and *tufA* mutants. PitA functions as a zinc uptake system when the zinc concentration is below the minimum inhibitory concentration (MIC) [14], RimP is required for 30S ribosome maturation [15], and Elongation factor Tu1 (*tufa*) is required for ribosomal peptide elongation [16].

**Table 1.**
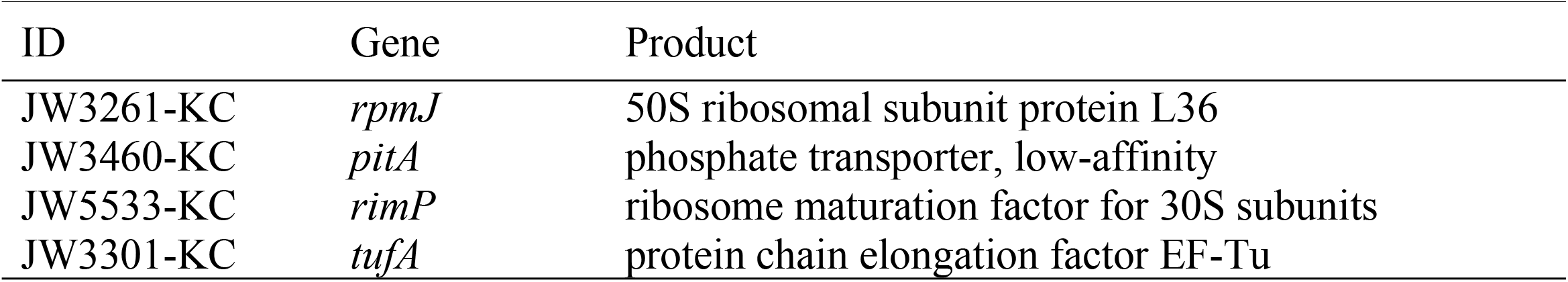
*E. coli* gene knockout mutans resistant to zinc.

We performed a complementation test to confirm that zinc resistance is caused by a lack of *rpmJ*. The results demonstrated that introducing the *rpmJ* gene into the *rpmJ* mutant reduced the zinc resistance (**Fig 1**). In contrast, zinc resistance was not reduced by introducing mutant *rpmJ* genes in which C27 or H33, important amino acids for the zinc-finger structure of RpmJ [13, 17], were replaced with serine (**Fig 1**). These results indicate that the loss of RpmJ function by destroying the zinc-finger structure leads to zinc resistance in *E. coli*.

**Fig 1.**
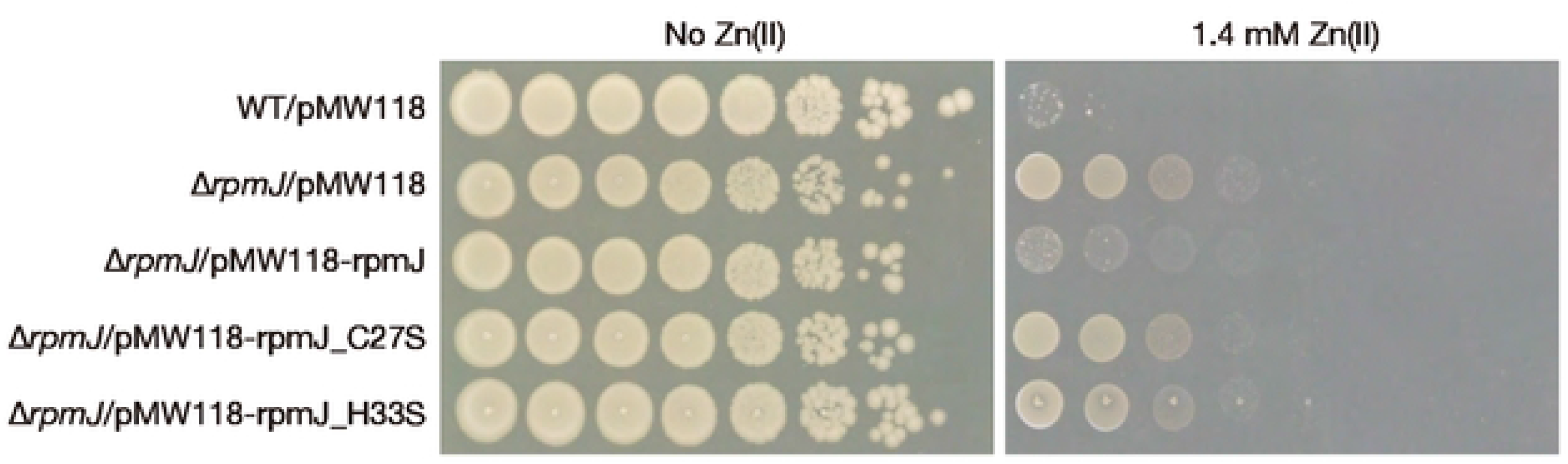
The *rpmJ* mutant exhibits zinc resistance. Overnight cultures of the wild-type strain transformed with an empty vector (WT/pMW118), the *rpmJ* mutants transformed with an empty vector (Δ*rpmJ*/pMW118), a plasmid carrying intact *rpmJ* gene (Δ*rpmJ*/pMW118-rpmJ), and plasmids carrying mutated *rpmJ* genes (Δ*rpmJ*/pMW118-rpmJ_C27S, Δ*rpmJ*/pMW118-rpmJ_H33S) were serially diluted 10-fold, spotted onto LB agar plates with or without 1.4 mM Zn(II), and incubated overnight at 37°C.

### Knockout of *rpmJ* alters ribosomal function

Given that RpmJ is a ribosomal protein, its knockout could alter the ribosomal structure. We examined the sensitivity of the *rpmJ* mutant to protein synthesis inhibitors that target ribosomes. Compared with the wild-type strain, the growth of the *rpmJ* mutant was decreased by all 4 tested inhibitors, chloramphenicol, erythromycin, clarithromycin, and tetracycline (**Fig 2**). This finding suggested that the *rpmJ* mutant has ribosomal structure changes by which protein synthesis inhibitors can easily access target sites of ribosomes. We considered that such structural changes could affect the translational function of the ribosome, and measured the translation fidelity using a dual luciferase assay in which stop codon readthroughs or frameshift readthroughs were detected (**Fig 3A**) [18]. In the assay, stop codons or frameshift mutations are inserted between Rluc and Fluc genes, and a low Fluc/Rluc (F/R) value indicates that the translation is accurate [18]. In the UGA stop codon readthrough, the F/R values were higher in the *rpmJ* mutant than in the wild-type strain in both the no-zinc and 0.8-mM zinc conditions (**Fig 3B**). In the UAG stop codon readthrough, the F/R values did not differ between the wild-type strain and the *rpmJ* mutant in the no-zinc and 0.8-mM zinc conditions (**Fig 3B**). In the +1 frameshift readthrough, the F/R value was higher in the *rpmJ* mutant than in the wild-type strain in the no-zinc condition, but no difference was observed in the 0.8-mM zinc condition (**Fig 3B**). In the −1 frameshift readthrough, the F/R value did not differ between the wild-type strain and the *rpmJ* mutant in the no-zinc condition, but the F/R value was lower in the *rpmJ* mutant than in the wild-type strain in the 0.8-mM zinc condition (**Fig 3B**). These results suggest that the ribosomal function required to maintain translation fidelity was altered in the *rpmJ* mutant.

**Fig 2.**
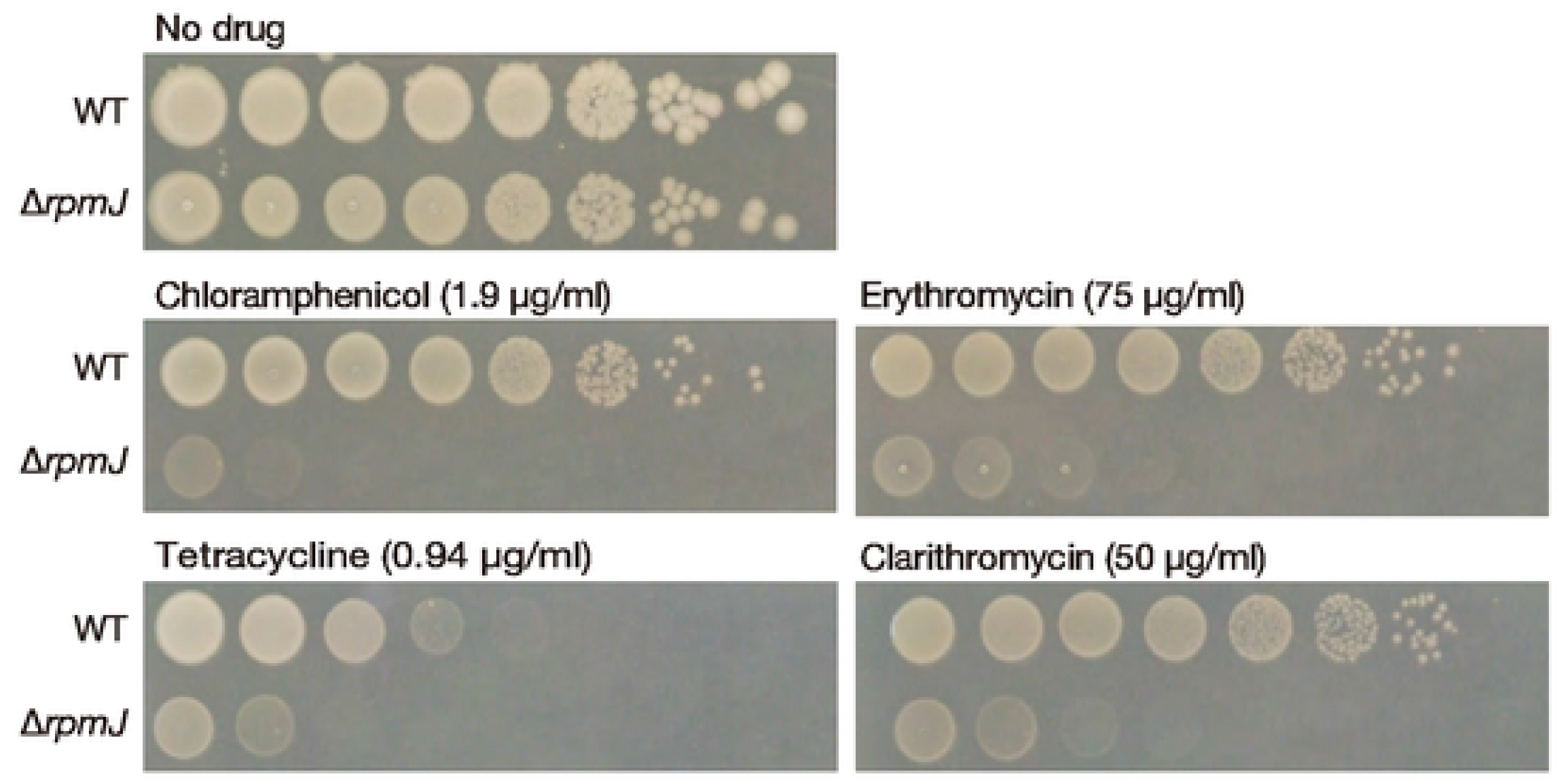
The *rpmJ* mutant is sensitive to protein synthesis inhibitors. Overnight cultures of the wild-type strain (WT) and the *rpmJ* mutant (Δ*rpmJ*) were serially diluted 10-fold, spotted onto LB agar plates with or without chloramphenicol (1.9 μg/ml), erythromycin (75 μg/ml), tetracycline (0.94 μg/ml), or clarithromycin (50 μg/ml), and incubated overnight at 37°C.

**Fig 3.**
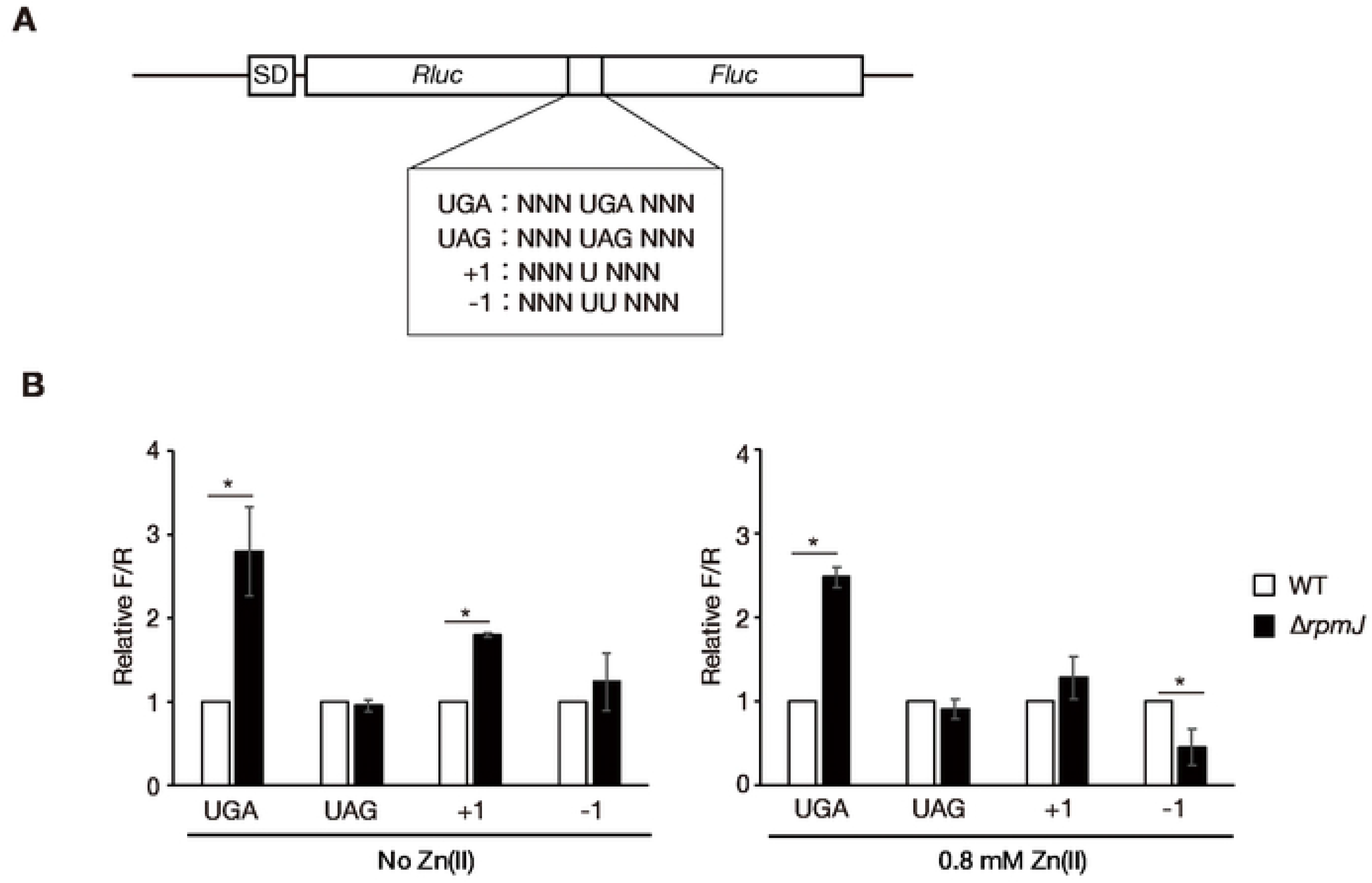
The *rpmJ* mutant had altered translational fidelity. A. The structure of the luciferase genes used for the dual-luciferase assay is shown. Stop codons or frameshift mutations are located between the Fluc and Rluc genes. Fluc and Rluc are expressed as a fusion protein when reading through stop codons or when misreading frameshift mutations occur.
B. The wild-type strain (WT) and the *rpmJ* mutant (Δ*rpmJ*) were cultured in the presence or absence of 0.8 mM Zn(II) and luciferase activity was measured. The F/R values normalized by that of the wild-type are indicated on the vertical axis. Data shown are means ± standard deviation from 3 independent experiments. The asterisk represents a p value <0.05.

### The *rpmJ* mutant has a low intracellular zinc concentration under excess zinc conditions

The ability of the *rpmJ* mutant to grow in an excess zinc condition could be due to a low intracellular zinc concentration. We measured the intracellular zinc concentration by inductively coupled plasma-mass spectrometry (ICP-MS) [19]. In a no-zinc and a 0.6-mM zinc conditions, the intracellular zinc concentrations did not differ between the wild-type strain and *rpmJ* mutant (**Fig 4**). In contrast, in a 1.2-mM excess zinc condition, the intracellular zinc concentration was lower in the *rpmJ* mutant than in the wild-type strain (**Fig 4**). In addition, in the 1.2-mM excess zinc condition, the intracellular zinc concentration did not differ between the wild-type strain and the *rpmJ* mutant transformed with the intact *rpmJ* gene (**Fig 4**). These results suggest that the *rpmJ* mutant maintained a low intracellular zinc concentration under an excess zinc condition, which could confer zinc resistance to the *rpmJ* mutant.

**Fig 4.**
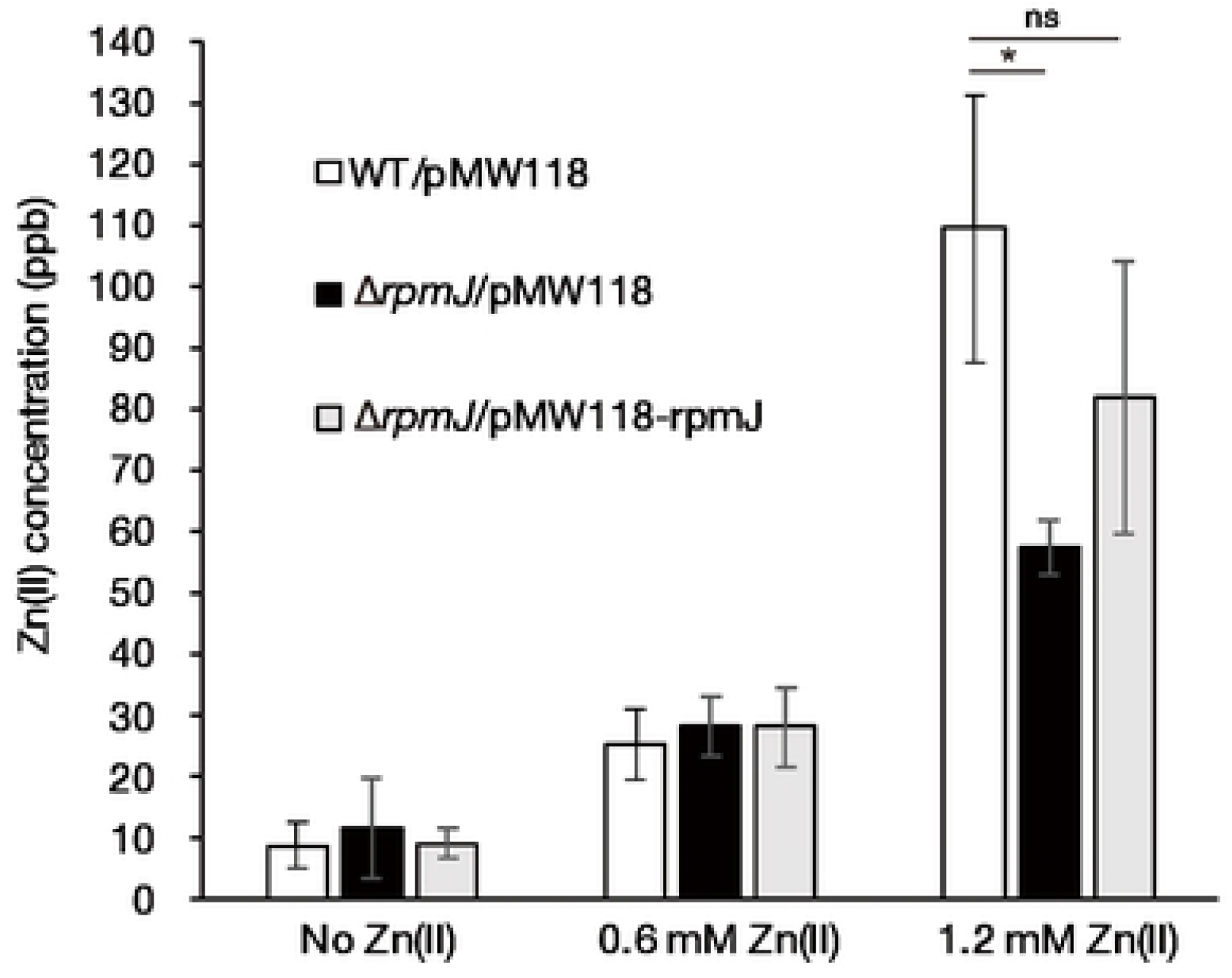
The intracellular zinc(II) concentration in the *rpmJ* mutant is low under excess zinc conditions. Wild-type *E. coli* strain transformed with an empty vector (WT/pMW118), the *rpmJ* mutant transformed with an empty vector (Δ*rpmJ*/pMW118), the *rpmJ* mutant transformed with a plasmid carrying an intact *rpmJ* gene (Δ*rpmJ*/pMW118-rpmJ) were cultured under conditions of 0 mM Zn(II), 0.6 mM Zn(II), and 1.2 mM Zn(II). The zinc concentration was measured by ICP-MS. Data shown are means ± standard deviation from 4 independent experiments. The asterisk represents a p value <0.05.

### Expression of iron-sulfur cluster synthesis genes is decreased in the *rpmJ* mutant

To understand the molecular mechanisms underlying the zinc resistance of the *rpmJ* mutant, we performed RNA sequence analysis to identify differentially expressed genes in the *rpmJ* mutant. In the *rpmJ* mutant, 195 genes were upregulated and 275 genes were downregulated compared with the wild-type strain (**S1 Table**). Contrary to our expectation, the expression of zinc uptake or zinc efflux genes was not altered in the *rpmJ* mutant. In contrast, expression of 6 genes encoding synthases for iron-sulfur clusters was decreased in the *rpmJ* mutant (**S1 Table**). Because iron-sulfur clusters are toxic targets of zinc, decreased expression of synthesis genes could decrease the amounts of iron-sulfur clusters and contribute to the zinc resistance of the *rpmJ* mutant. To elucidate characteristic features of the differentially expressed genes in the *rpmJ* mutant, we performed a gene ontology (GO) enrichment analysis. The upregulated genes included those categorized as related to translation or ribosomal subunits (**Fig 5**), suggesting that ribosomal function is damaged in the *rpmJ* knockout and some compensatory regulatory mechanisms were triggered to increase translation function.

**Fig 5.**
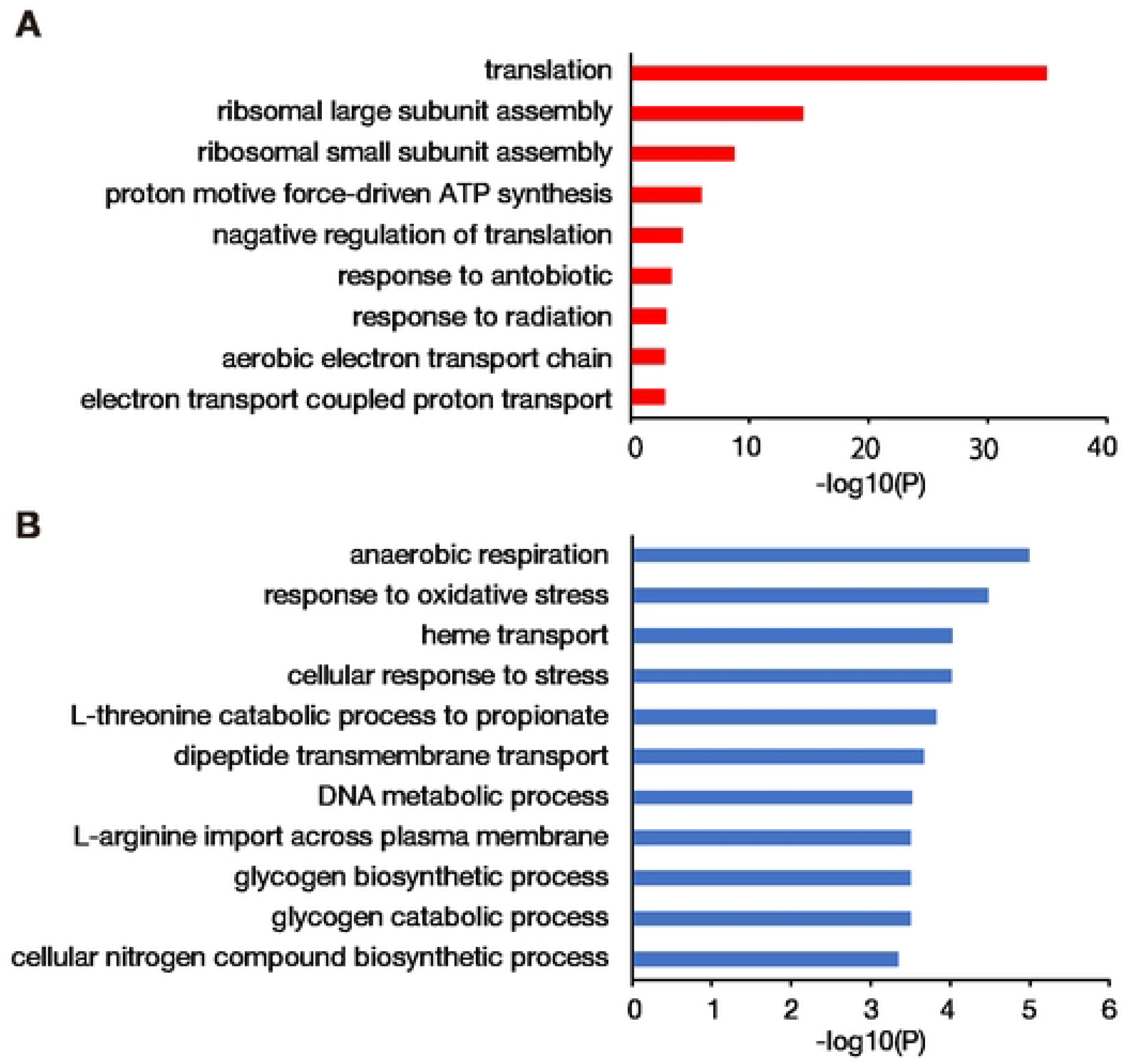
GO enrichment analysis of differentially expressed genes in the *rpmJ* mutant. GO enrichment analysis was performed using differentially expressed genes in the *rpmJ* mutant (195 upregulated genes, 275 downregulated genes) (**S1 Table**). Categories with a p value <0.001 are shown. GO-enriched categories of genes with increased expression are shown in panel A and categories of genes with decreased expression are shown in panel B.

### Knockout of several ribosomal proteins leads to a zinc resistance phenotype

*E. coli* has 7 nonessential ribosomal proteins other than RpmJ. We examined whether knockout of these nonessential ribosomal proteins leads to zinc resistance as in the case of the *rpmJ* knockout. Knockout of *rplA*, *rpmE*, *rpmI*, and *rpsT* also caused zinc resistance (**Fig 6**). The results suggest the existence of some conserved zinc resistance mechanisms among the gene knockout mutants of ribosomal proteins.

**Fig 6.**
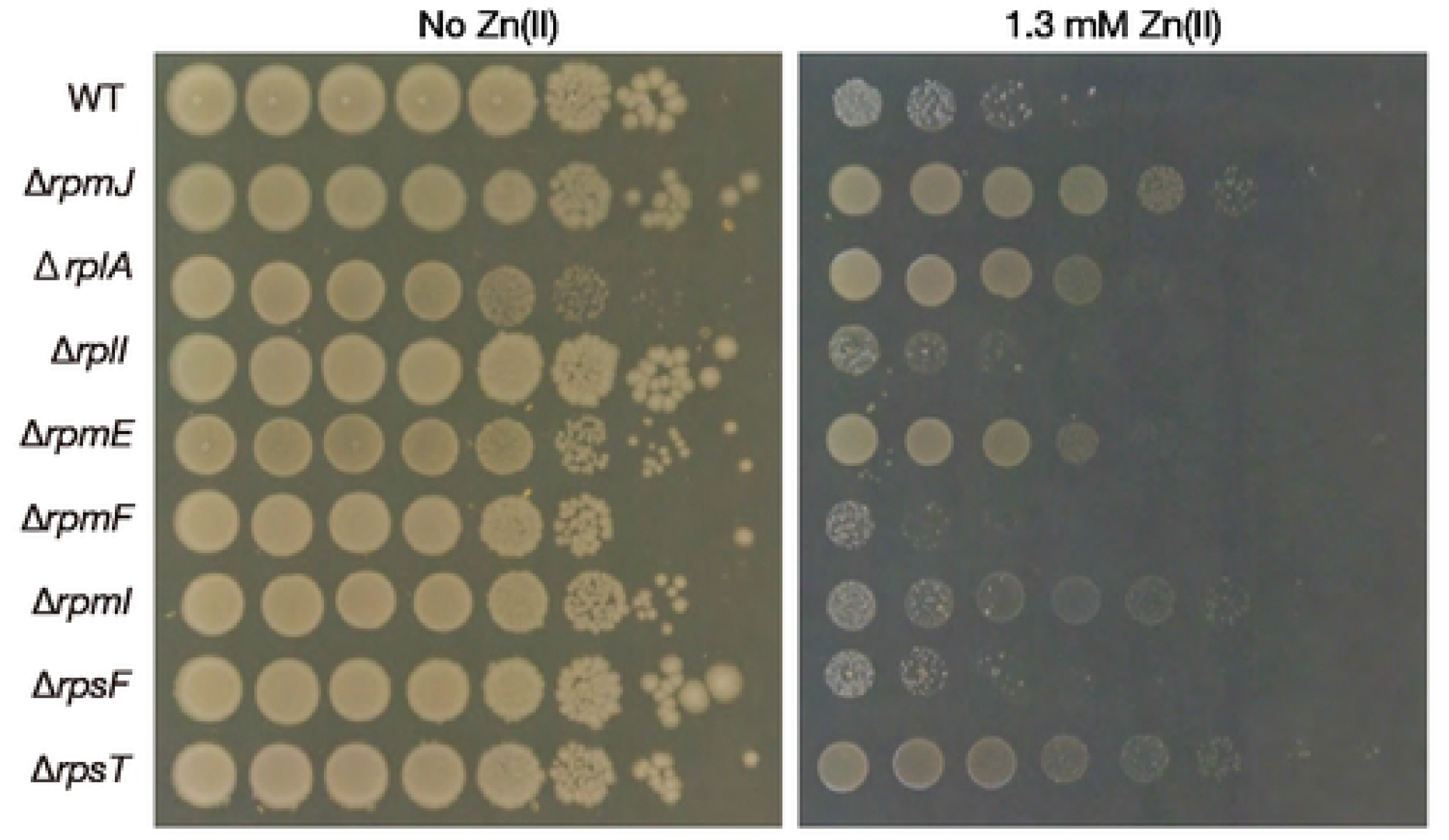
Knockout mutants of nonessential ribosomal proteins exhibit zinc resistance. Overnight cultures of the wild-type strain and knockout mutants of ribosomal proteins (RpmJ, RplA, RplI, RpmE, RpmF, RpmI, RpsF, and RpsT) were serially diluted 10-fold, spotted onto LB agar plates with or without 1.3 mM Zn(II), and incubated overnight at 37°C.

## Discussion

The present findings revealed that knocking out ribosomal protein RpmJ confers zinc resistance to *E. coli*. The *rpmJ* mutant had a low concentration of intracellular zinc and decreased expression of iron-sulfur cluster synthesis genes. In addition, knocking out other ribosomal proteins, including RplA, RpmE, RpmI, and RpsT, led to zinc resistance in *E. coli*. This study is the first to reveal that ribosomal protein deficiency causes *E. coli* resistance to zinc.

Mechanisms of zinc resistance or sensitivity have been investigated by identifying mutations of zinc uptake or efflux systems [20]. In the present study, we identified novel gene mutations other than those of transporters that confer zinc resistance. RNA sequence analysis did not reveal differentially expressed genes involved in zinc efflux or uptake. Because the RNA sequence analysis used RNA samples prepared under a no-zinc condition, it is possible that zinc efflux or uptake genes were differentially expressed in the *rpmJ* mutant under excess zinc conditions. In addition, RNA sequence analysis identified that the *rpmJ* mutant had decreased expression of genes involved in the synthesis of iron-sulfur clusters. The lower intracellular zinc concentration and downregulated expression of iron-sulfur cluster synthesis genes are plausible molecular mechanisms for the zinc resistance of the *rpmJ* mutant.

The *rpmJ* mutant was sensitive to protein synthesis inhibitors, and exhibited altered translation fidelity and increased expression of ribosomal subunit genes. RNA sequence analysis also revealed altered expression of many genes other than ribosome-related genes in the *rpmJ* mutant. These findings suggest that structural abnormalities or functional alterations of ribosomes in the *rpmJ* mutant are sensed by some transcriptional regulators, leading to differential transcription of various genes. Ribosomal proteins are able to repress their own gene translation [21], but the effects on other genes are not known. The stringent response is a well-known phenomenon that regulates the transcription of many genes when amino acids are limited and translation is inhibited [22]. In the stringent response, tRNA without an amino acid enters into the ribosome A-site and activates RelA protein, a synthase of ppGpp. ppGpp produced by RelA activates the transcription of various genes [23, 24]. The altered structure or dysfunction of ribosomes in the *rpmJ* mutant may result in activation of RelA to induce the expression of various genes. This point should be investigated in future studies.

Previous studies demonstrated that 8 ribosomal proteins interact with zinc [25, 26]. Among the 5 ribosomal proteins whose knockout leads to zinc resistance, RpmJ and RpmE interact with zinc [27]. Under zinc-limited conditions, RpmJ and RpmE are released from ribosomes and supply zinc by self-degradation [28–32]. In contrast, RplA, RpmI, and RpsT, whose knockout leads to zinc resistance, do not interact with zinc and do not function in zinc homeostasis. Thus, the capacity of the ribosomal protein to interact with zinc is not related to the zinc resistance conferred by the knockout of the ribosomal protein. We speculate that some abnormalities of the ribosomal structure and function are conserved among the ribosomal protein mutants that showed zinc resistance. The present study also demonstrated that knockout of *rimP*, involved in 30S ribosome maturation [15], and *tufA*, involved in ribosomal peptide elongation [16], leads to zinc resistance in *E. coli*. The *rimP-* and *tufA-* knockout mutants could have ribosomal abnormalities and may have the same zinc-resistant mechanisms as the ribosomal protein mutants. Further studies are needed to clarify the molecular mechanisms underlying zinc resistance by investigating ribosomal structure and function in the zinc-resistant mutants identified in this study.

## Materials and methods

### Bacterial strains and culture conditions

*E. coli* BW25113 and the gene knockout strains were cultured on LB agar medium, and the colonies were aerobically cultured in LB liquid medium at 37°C. *E. coli* harboring pMW118 was cultured on LB agar plates containing 100 μg/ml ampicillin. The bacterial strains and plasmids used in this study are listed in **Table 2**.

**Table 2.** List of bacterial strains and plasmids used.

| Strain or plasmid | Genotypes or characteristics | Source or reference |
| --- | --- | --- |
| **Strains** |  |  |
| BW25113 | *rrnB*, Δ*lacZ*4787, *HsdR*514, $\Delta$(*araBAD*)567, $\Delta$(*rhaBAD*)568, *rph*-1 | NBRP |
| JW3261-KC | BW25113 Δ*rpmJ*::*kan* Kan<sup>r</sup> | NBRP |
| JW3947-KC | BW25113 Δ*rplA*::*kan* Kan<sup>r</sup> | NBRP |
| JW4161-KC | BW25113 Δ*rplI*::*kan* Kan<sup>r</sup> | NBRP |
| JW3907-KC | BW25113 Δ*rpmE*::*kan* Kan<sup>r</sup> | NBRP |
| JW1075-KC | BW25113 Δ*rpmF*::*kan* Kan<sup>r</sup> | NBRP |
| JW1707-KC | BW25113 Δ*rpmI*::*kan* Kan<sup>r</sup> | NBRP |
| JW4158-KC | BW25113 Δ*rpsF*::*kan* Kan<sup>r</sup> | NBRP |
| JW0022-KC | BW25113 Δ*rpsT*::*kan* Kan<sup>r</sup> | NBRP |
| JW3460-KC | BW25113 $\Delta$ *pitA*::*kan* Kan<sup>r</sup> | NBRP |
| JW5533-KC | BW25113 Δ*rimP*::*kan* Kan<sup>r</sup> | NBRP |
| JW3301-KC | BW25113 $\Delta$ *tufA*::*kan* Kan<sup>r</sup> | NBRP |
| JM109 | Host strain for cloning | Takara Bio |
| **Plasmids** |  |  |
| pMW118 | Low-copy-number plasmid; Amp<sup>r</sup> | Nippon Gene |
| pMW118-rpmJ | pMW118 with *rpmJ*; Amp<sup>r</sup> | This study |
| pMW118-rpmJ_C27S | pMW118 with C27S *rpmJ*; Amp<sup>r</sup> | This study |
| pMW118-rpmJ_H33S | pMW118 with H33S *rpmJ*; Amp<sup>r</sup> | This study |
| pQE-Luc(UGA) | pQE60 with UGA window between Fluc and Rluc; Amp<sup>r</sup> | [18] |
| pQE-Luc(UAG) | pQE60 with UAG window between Fluc and Rluc; Amp<sup>r</sup> | [18] |
| pQE-Luc(+1) | pQE60 with +1 window between Fluc and Rluc; Amp<sup>r</sup> | [18] |
| pQE-Luc(-1) | pQE60 with -1 window between Fluc and Rluc; Amp<sup>r</sup> | [18] |

Kan: kanamycin, Amp: ampicillin.

### Evaluation of bacterial resistance to antimicrobial substances

To measure bacterial resistance to zinc and antibiotics, autoclaved LB agar medium was mixed with ZnSO_4_•7H_2_O (Nacalai Tesque, Kyoto, Japan) or antibiotics and poured into square plastic dishes (Eiken Chemical, Tokyo, Japan). *E. coli* overnight cultures were serially diluted 10-fold in 96-well microplates, and 5 μl of the diluted bacterial solution was spotted onto the LB agar plates supplemented with drugs. The plates were incubated at 37°C for 1 day and colonies were photographed using a digital camera.

### Genetic manipulation

Gene knockout mutants were constructed by phage transduction using phage P1 *vir* from the gene knockout mutants in the Keio collection as donor strains to the BW25113 strain as the recipient strain (**Table 2**). To construct a plasmid carrying the *rpmJ* gene, a DNA fragment encoding the *rpmJ* gene was amplified by polymerase chain reaction (PCR) using primer pairs (rpmJ_F_XbaI_2nd and rpmJ_R_HindIII_2nd; **Table 3**) from genomic DNA of the BW25113 strain as a template. The amplified DNA fragment was cloned into XbaI and HindIII sites of pMW118, resulting in pMW118-rpmJ. Amino acid substitution mutations were introduced into pMW118-rpmJ by PCR using primer pairs (rpmj_C27S_F and rpmj_C27S_R or rpmj_H33S_F and rpmj_H33S_R; **Table 3**) and pMW118-rpmJ as a template. Mutations were confirmed by DNA sequencing.

**Table 3.**
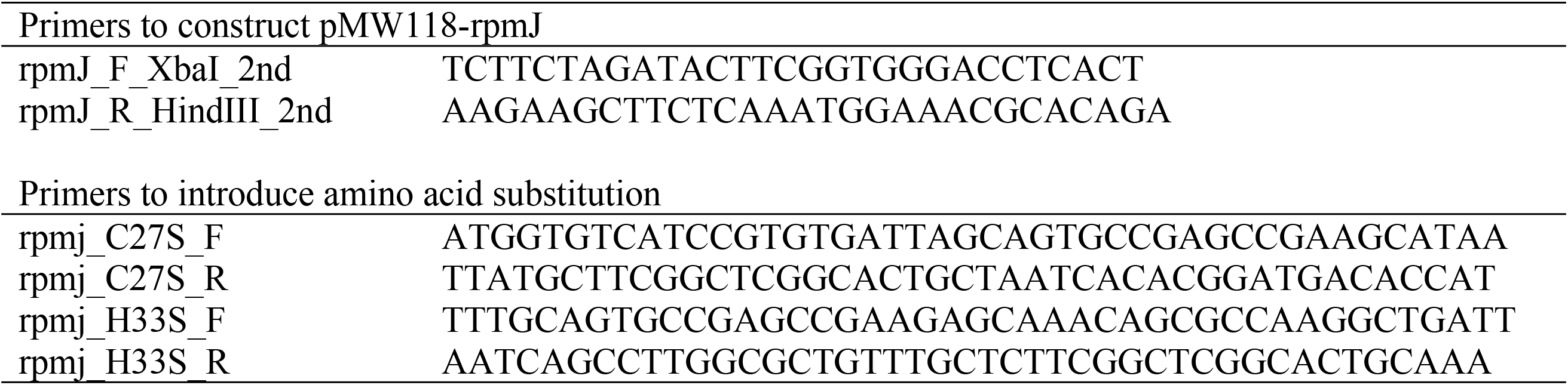
Primers used in this study.

### Dual-luciferase assay

The wild-type *E. coli* strain and *rpmJ* knockout mutant were transformed with plasmids [pQE-Luc(UGA), pQE-Luc(UAG), pQE-Luc(+1), pQE-Luc(−1)] [18] (**Table 2**). Each transformant was aerobically cultured in LB liquid medium containing 100 μg/ml ampicillin at 37°C overnight. The overnight culture was inoculated into a 100-fold amount of fresh LB medium. For cells in the no-zinc condition, cells were cultured until OD_600_=0.5 and then collected. For cells in the zinc condition, cells were cultured until OD_600_=0.25-0.35 in the no-zinc condition, supplemented with 0.8 mM Zn(II), and then further cultured for 1 h before collecting. The cell pellets were suspended in 200 μl buffer (50 mM HEPES-KOH [pH7.6], 100 mM KCl, 10 mM MgCl_2_, 7 mM β-mercaptoethanol, 400 μg/ml lysozyme). The cell sample was then subjected to freezing and thawing using liquid nitrogen and centrifuged at 15,000 rpm for 15 min at 4°C. The centrifuge supernatant was mixed with an equal volume of Firefly luciferase substrate (Promega) or Renilla luciferase substrate (Pierce), and the luminescence intensity was measured with a luminometer (Promega).

### Measurement of intracellular zinc concentration

Zinc concentrations were measured according to a previously reported method [19]. Briefly, 100 μl of *E. coli* overnight cultures were spread on agar plates supplemented with no zinc, 0.6 mM Zn(II), or 1.2 mM Zn(II), and cultured overnight at 37°C. The cells were suspended in phosphate buffered saline and the OD_600_ value was adjusted to 0.5. The sample was centrifuged, the bacterial pellet was washed 5 times with cold phosphate buffered saline, and 100 μl of 50% HNO_3_ was added. The sample was heated at 65°C overnight. The HNO_3_ concentration was adjusted to 5% and the zinc concentration was determined by ICP-MS (Agilent7500cx, Agilent Technologies).

### RNA-Sequence analysis

Total RNA of *E. coli* was extracted according to a previously described method [33] with minor modifications. *E. coli* overnight culture (50 μl) was inoculated into 5 ml LB medium and aerobically cultured at 37°C. When the OD_600_ of the culture reached 0.7, 1.8 ml of culture was vortex-mixed with 200 μl of 5% phenol in ethanol, chilled in ice water for 5 min, and centrifuged at 21,500×g for 2 min. The bacterial precipitate was frozen in liquid nitrogen and stored at −80°C for 2 h. The precipitate was dissolved in 200 μl lysis buffer (TE buffer, 1% lysozyme, 1% sodium dodecyl sulfate) and incubated at 65°C for 2 min. The sample was subjected to RNA extraction using an RNeasy minikit (Qiagen) according to the manufacturer’s protocol. rRNA was removed from the total RNA using a NEBNext rRNA depletion kit (NEB), and RNA was converted to a DNA library using a TruSeq stranded total RNA kit (Illumina). RNA sequencing was performed using a NovaSeq 6000 system (Illumina), and at least 4 billion base sequences of 100-base paired-end reads were generated per sample. The data were analyzed using CLC Genomics Workbench software (version 11.0). The reads were mapped to a reference genome of the *E. coli* W3110 strain (NCBI reference sequence NC_007779.1), and the reads per kilobase of transcript per million mapped reads (RPKM) were compared between the wild-type strain and the *rpmJ* mutant. The experiment was independently performed twice to identify the genes for which the mean values differed by >2-fold between BW25113 and Δ*rpmJ* and the false discovery rate p value was <0.001. GO analysis was performed using software developed by the European Molecular Biology Laboratory (https://www.ebi.ac.uk/QuickGO).

### Statistical analysis

Differences in dual luciferase assay were evaluated by Student’s *t* test in Excel. Differences in the intracellular zinc concentration by ICP-MS were evaluated by Dunnett’s test in GraphPad PRISM software.

## Supporting information

**S1 Table. Differentially expressed genes in the *rpmJ* knockout mutant**.

Yellow background indicates iron-sulfur cluster synthesis genes.

## Acknowledgement

We thank the Okayama University Institute of Plant Science and Resources, for their help in ICP-MS measurements, and the National BioResource Project-*E. coli* (National Institute of Genetics, Japan) for providing the Keio collection.

